# Phylogenetic distribution of roseobacticides in the *Roseobacter* group and their effect on microalgae

**DOI:** 10.1101/242842

**Authors:** Eva C. Sonnenschein, Christopher Broughton William Phippen, Mikkel Bentzon-Tilia, Silas Anselm Rasmussen, Kristian Fog Nielsen, Lone Gram

## Abstract

The *Roseobacter-group* species *Phaeobacter inhibens* produces the antibacterial tropodithietic acid (TDA) and the algaecidal roseobacticides with both compound classes sharing part of the same biosynthetic pathway. The purpose of this study was to investigate the production of roseobacticides more broadly in TDA-producing roseobacters and to compare the effect of producers and non-producers on microalgae. Of 33 roseobacters analyzed, roseobacticide production was a unique feature of TDA-producing *P. inhibens, P. gallaeciensis* and *P. piscinae* strains. One TDA-producing *Phaeobacter* strain, 27-4, was unable to produce roseobacticides, possibly due to a transposable element. TDA-producing *Ruegeria mobilis* and *Pseudovibrio* did not produce roseobacticides. Addition of roseobacticide-containing bacterial extracts affected the growth of the microalgae *Rhodomonas salina, Thalassiosira pseudonana* and *Emiliania huxleyi*, while growth of *Tetraselmis suecica* was unaffected. During co-cultivation, growth of *E. huxleyi* was initially stimulated by the roseobacticide producer DSM 17395, while the subsequent decline in algal cell numbers during senescence was enhanced. Strain 27-4 that does not produce roseobacticides had no effect on algal growth. Both bacterial strains, DSM 17395 and 27-4, grew during co-cultivation presumably utilizing algal exudates. Furthermore, TDA-producing roseobacters have potential as probiotics in marine larviculture and it is promising that the live feed *Tetraselmis* was unaffected by roseobacticides-containing extracts.

**Originality-significance statement:** Some *Roseobacter-group* bacteria produce the antibacterial compound tropodithetic acid (TDA) and have potential as probiotics in marine aquaculture. However, a few of these strains additionally produce algaecidal compounds, the roseobacticides, which would restrict their use in marine larviculture where algae are used as live feed for fish larvae. We herein found that roseobacticides are limited to TDA-producing *Phaeobacter* strains and were not biosynthesized by TDA-producers outside this genus. Roseobacticides affected several strains of microalgae, but not the chlorophyte that is used as live feed in the aquaculture industry. Thus, the application of *Roseobacter* strains as probiotics is not hampered. Furthermore, these results demonstrate how *Roseobacter-group* strains act as gardeners of microalgae and thereby would be involved in environmental processes on a larger scale.

## Introduction

Microalgae are responsible for half of the global primary production and are the basis of the marine food web (Field *et al*., 1998). The phycosphere surrounding an algal cell is inhabited by bacteria that benefit from increased nutrient concentrations (Cole, 1982; Azam and Malfatti, 2007; Amin *et al*., 2012). In return, these bacteria may provide the algae with supplements such as vitamins (Croft *et al*., 2005; Cooper and Smith, 2015) and protect them from infections or grazing (Seyedsayamdost, Case, *et al*., 2011). The bacteria may, however, also act as pathogens or parasites causing lysis and death of the algal cells (Seyedsayamdost, Case, *et al*., 2011; Riclea *et al*., 2012; Wang *et al*., 2014). Possibly, these modes are interlinked genetically, or metabolically, and depend on exterior factors such as nutrient supply. This scenario was proposed for the interaction between *Emiliania huxleyi*, a globally important, bloom-forming haptophyte (Read *et al*., 2013) and the alphaproteobacterium *Phaeobacter inhibens* (Seyedsayamdost, Case, *et al*., 2011; Segev *et al*., 2016b; Wang *et al*., 2016; Wang and Seyedsayamdost, 2017). The bacterium belongs to the *Roseobacter* group, one of the most common and widespread marine bacterial groups and a group often co-appearing with microalgae (Sapp *et al*., 2007; Luo and Moran, 2014; Tan *et al*., 2015; Eva C Sonnenschein *et al*., 2017). *Roseobacter-group* bacteria can be involved in the degradation of the algae-produced dimethylsulfoniopropionate into dimethylsulfide, which is released into the atmosphere and serves as cloud nuclei (Miller and Belas, 2004; Dickschat *et al*., 2010). *P. inhibens* produces at least two groups of bioactive molecules: the antibacterial compound tropodithietic acid (TDA), and the algaecidal roseobacticides, the production of the latter being induced by p-coumaric acid, an algal degradation product (Brinkhoff *et al*., 2004; Seyedsayamdost, Carr, *et al*., 2011). First identified as its tautomer thiotropocin in 1984, TDA is a small acid comprising sulfur and a 7-membered carbon ring (Kintaka *et al*., 1984; Brock *et al*., 2013; D’Alvise *et al*., 2016). The roseobacticide family was discovered in 2011 and contains roseobacticide A to K with roseobacticide A and B generally being the dominant forms (Seyedsayamdost, Carr, *et al*., 2011; Seyedsayamdost, Case, *et al*., 2011). Like TDA, the roseobacticides also contain at least one sulfur atom and a tropone ring. Furthermore, they contain additional aromatic side chains derived from aromatic amino acids. It has been hypothesized that under nutrient-rich conditions, *P. inhibens* produces TDA to protect *E. huxleyi* from pathogens and, when *E. huxleyi* senesces, the algal degradation product would induce production of roseobacticides in *P. inhibens*, enhancing the algal decay. While the metabolic pathways of both compound groups are not fully elucidated (Geng *et al*., 2008; Brock *et al*., 2014; Seyedsayamdost *et al*., 2014; Wang *et al*., 2016), it has been shown for *P. inhibens* DSM 17395 that the production of TDA and roseobacticides are interlinked, i.e. that the same genes are essential for the biosynthesis of both compounds and that the metabolites probably share the same precursor (Wang *et al*., 2016).

TDA-producing roseobacters are also of interest in biotechnological application as potential probiotic bacteria in marine aquaculture (Bruhn *et al*., 2005; D’Alvise *et al*., 2012, 2013; Bentzon-Tilia *et al*., 2016). Especially, fish larvae are prone to bacterial infections due to their undeveloped immune system and probiotics are a promising alternative to the use of antibiotics that can result in development and spread of antibiotic resistance (Cabello, 2006). The fish larvae are fed with live feed such as rotifers and *Artemia* that are themselves fed with live microalgae. Pathogenic bacteria may proliferate in these feed cultures due to high levels of nutrients (Verdonck *et al*., 1997). *P. inhibens* is an excellent antagonist of fish pathogenic Vibrionaceae due to the production of TDA and can kill or inhibit fish pathogens in the live feed and reduce mortality caused by vibriosis in fish larvae (D’Alvise *et al*., 2012, 2013; Grotkjær *et al*., 2016). So far, no negative effect of *P. inhibens* was found on the aquaculture organisms themselves (Neu *et al*., 2014), but pure TDA was able to change the natural microbiota of the feed algae *Nannochloropsis salina* (Geng *et al*., 2016). However, it is obviously of concern that the bacterium potentially can harm the microalgae used for feeding.

The aim of this study was to analyze the distribution of roseobacticides within the *Roseobacter* group and determine which TDA-producing organisms also produce roseobacticides. We compared the genomes of producers and non-producers to identify genes likely contributing to the biosynthesis of roseobacticides. Furthermore, we investigated the impact of a roseobacticide producer and a non-producer on microalgae using bacterial extracts as well as co-cultivation.

## Results and discussion

### Phylogenetic distribution of roseobacticides

Thirty-three *Roseobacter-group* bacteria, including 27 TDA producers, were analyzed for the ability to biosynthesize roseobacticides upon induction with p-coumaric acid (see Supplementary materials & methods for details). If the dominant roseobacticide, roseobacticide B, was detected in the ethyl acetate extract of the culture by mass spectrometry, the strain was considered positive for production of roseobacticides (Table 1, Fig. S1, Table S1). With the exception of *P. piscinae* 27-4 and *P. porticola* P97, all tested *Phaeobacter* strains produced both, TDA and roseobacticide B (Fig. 1). Interestingly, the roseobacticide producer 8-1 was even collected in the same location and year as 27-4 underlining the high phenotypic diversity among genetically similar *Roseobacter-group* strains (Table 1) (Sonnenschein *et al*., 2017). Strains from the closely related genera *Pseudophaeobacter* and *Leisingera* as well as TDA-producers from the genera *Ruegeria* and *Pseudovibrio* did not produce roseobacticides under the tested conditions. This confirmed previous roseobacticide analyses of the strains DSM 17395 (BS107), 2.10, 27-4 and TM1040, and adds DSM 16374 as a roseobacticide producer although not detected previously (Seyedsayamdost, Carr, *et al*., 2011; Wang and Seyedsayamdost, 2017). Thus, the ability of roseobacticide production appears to be limited to the phylogenetic neighbours *P. inhibens, P. gallaeciensis* and *P. piscinae*. In contrast, TDA is produced by several *Roseobacter-group* bacteria, which are not closely related. Evolution of the TDA genes is in agreement with the phylogenetic clustering of the strains (Fig. S2). We propose that a common ancestor of *P. inhibens, P. gallaeciensis* and *P. piscinae* obtained the TDA biosynthetic genes e.g. by horizontal gene transfer and subsequently, additionally developed the ability to produce roseobacticides. This hypothesis is support by the findings by Wang *et al*., who demonstrated a tight genetic and metabolic dependency of roseobacticides to TDA in *P. inhibens* DSM 17395 (Wang *et al*., 2016).

**Table 1.**
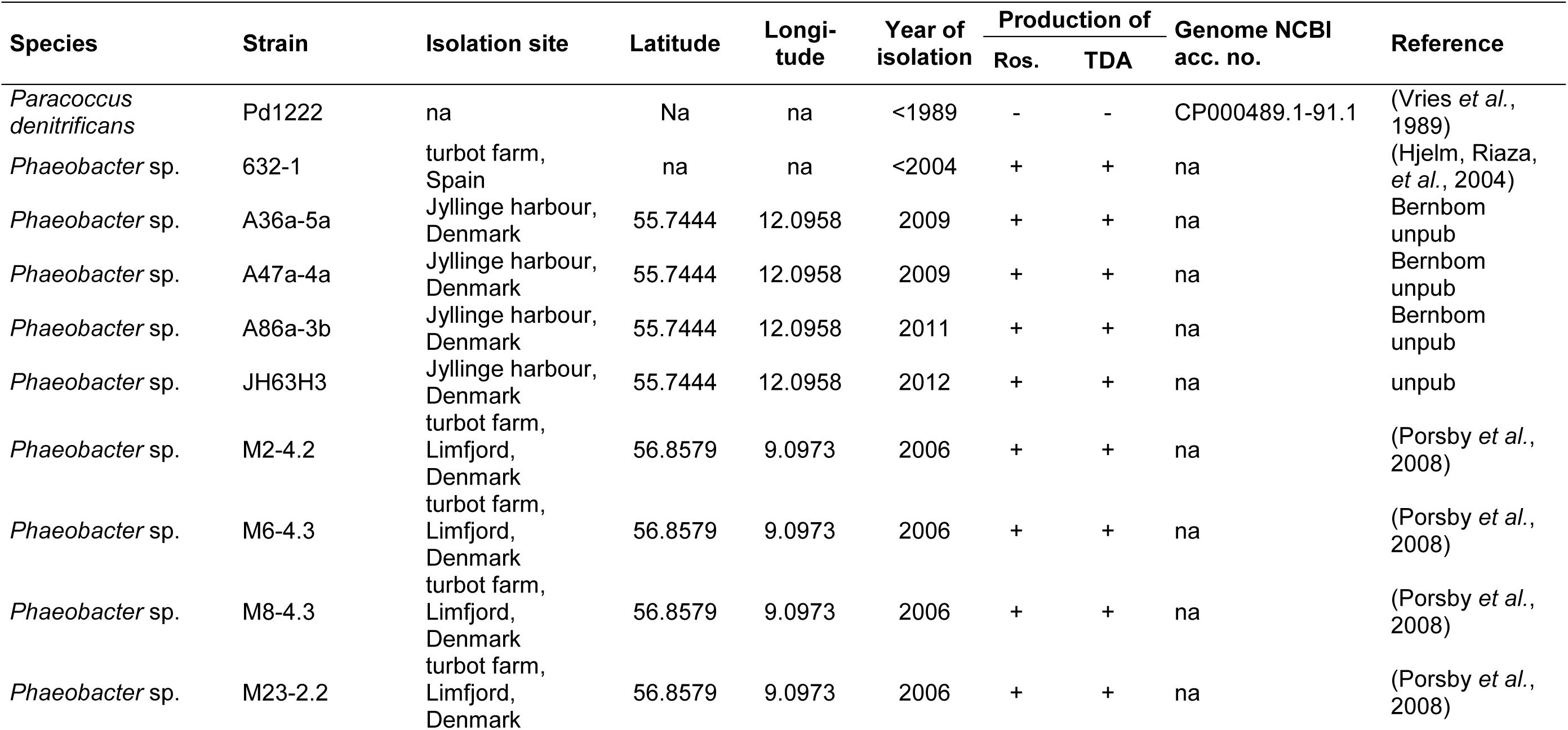
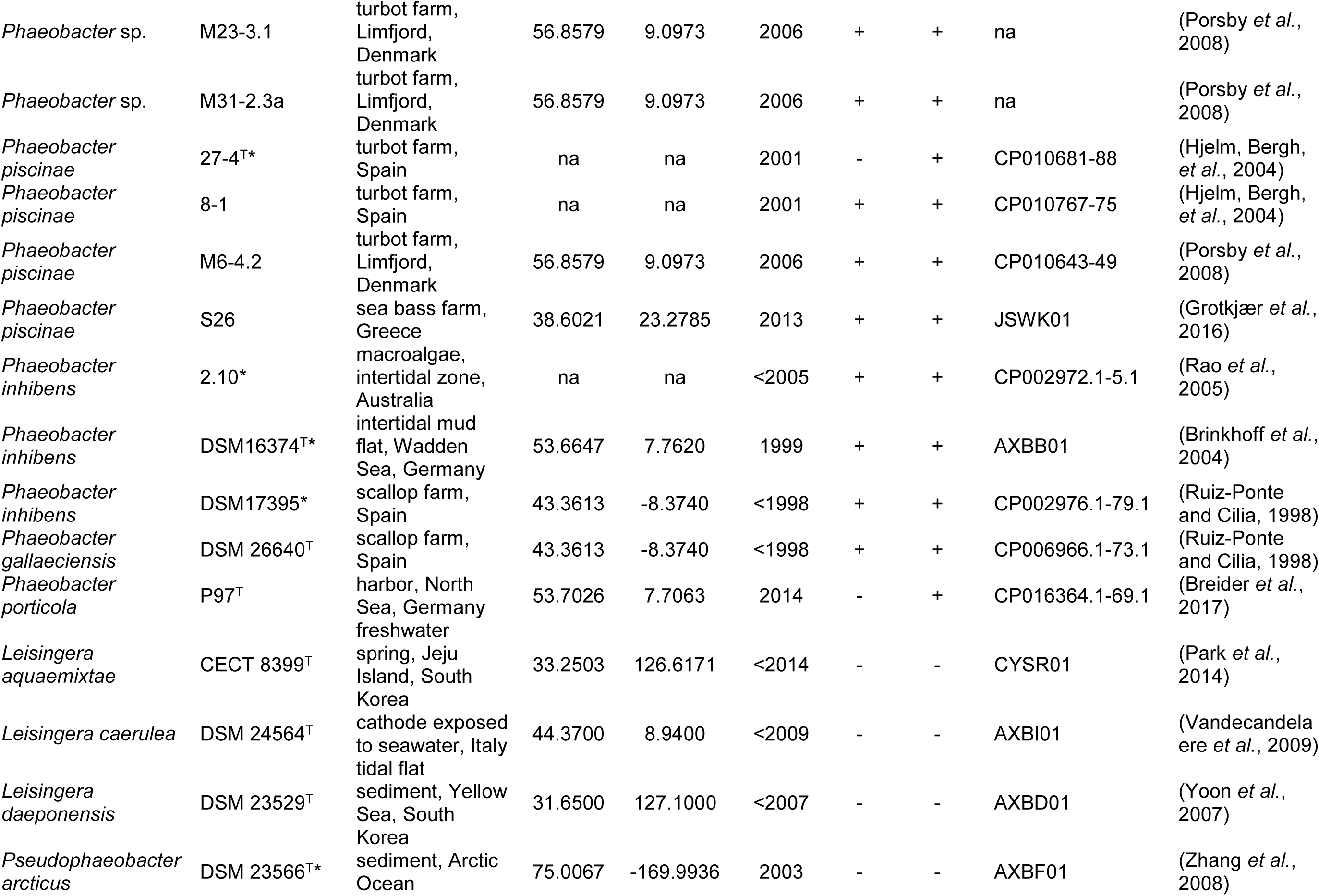
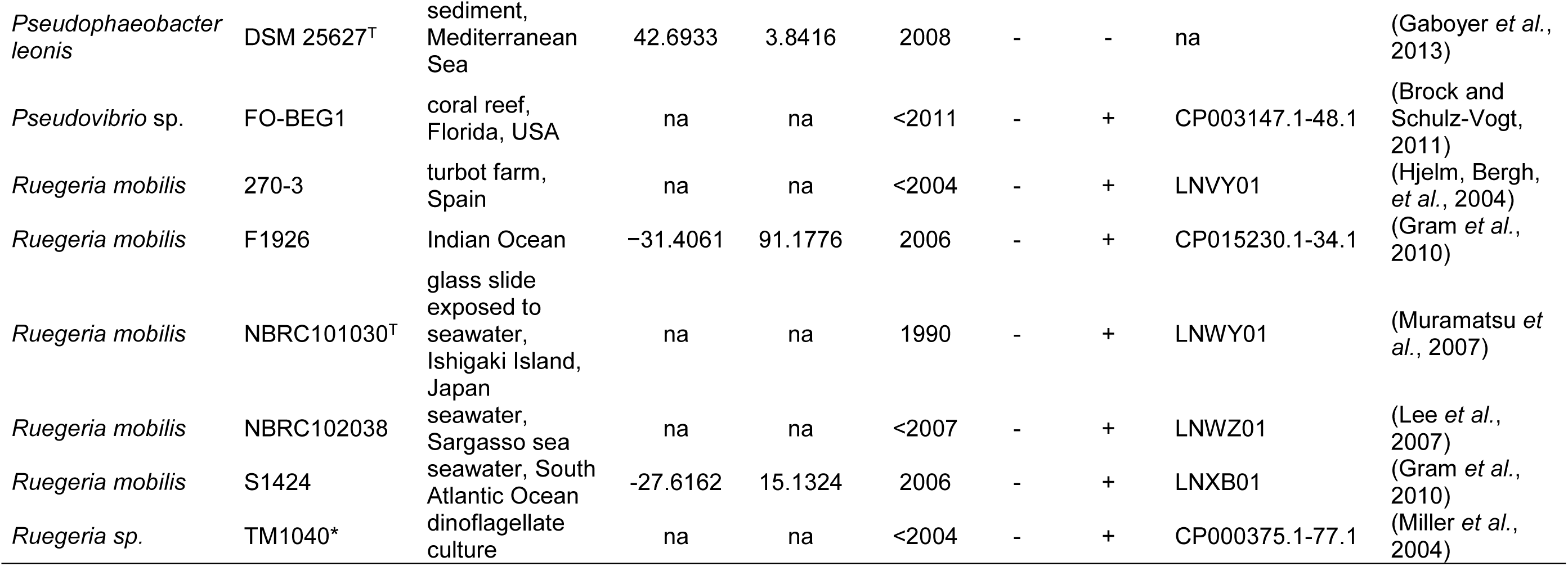
Strains evaluated for roseobacticide and TDA production in this study. Ros. = roseobacticides, TDA = tropodithietic acid; na = not available; unpub = unpublished. * = strains that were also previously analyzed for roseobacticide production (Seyedsayamdost, Carr, *et al*., 2011; Wang and Seyedsayamdost, 2017)

**Figure 1.**
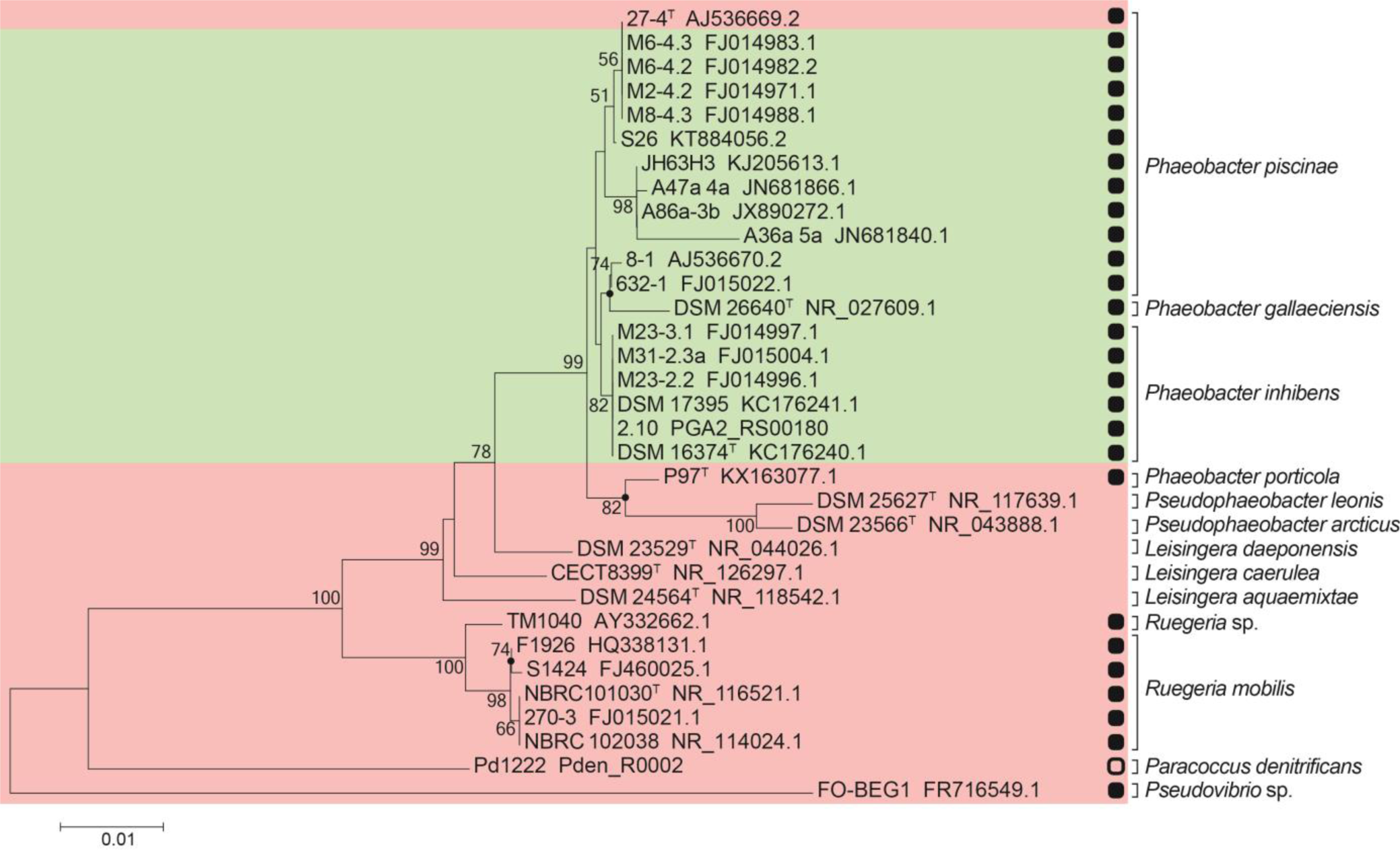
Neighbor-joining (NJ) tree of 16S rRNA gene sequences of *Roseobacter* group strains tested for production of roseobacticide B. The NJ analysis was performed using the bootstrap test with 1000 replicates. Support values (percentages) from 1000 bootstrap replicates are displayed above branches. Only values >50% are shown at major nodes. Filled circles indicate nodes also recovered reproducibly with the maximum-likelihood method. Roseobacticide B producers are marked in green, non-producers in red. TDA-producers are marked with a black square and the strain carrying the disrupted TDA biosynthetic gene cluster with an empty square.

P. *piscinae* 27-4 produced TDA, while roseobacticides were not detected. The species *P. piscinae* is phylogenetically distinct from the species *P. inhibens* and *P. gallaeciensis* (Sonnenschein *et al*., 2017), but based on whole genome comparison, 27-4 is very similar to at least three other *P. piscinae* strains (average nucleotide identity (ANI) to M6-4.2, S26, and 8-1 is 98.5, 98.3, 95.9%, respectively). To investigate the genetic background of roseobacticide production, the predicted protein sequences of the roseobacticide producer strains M6-4.2, 8-1, DSM 17395 and DSM 26640 and the non-producer 27-4 were compared using OrthoVenn (Wang *et al*., 2015). This revealed fourteen orthologous protein clusters unique to the producer strains (Fig. S3, see ^a^; Table 2A) and five unique to the non-producer (Fig. S3, see ^b^; Table 2B). The proteins unique to the producer strains differ from those found by Wang *et al*. (Wang *et al*., 2016) as being essential for the production of roseobacticides in DSM 17395 using a transposon library; however, some lie within close proximity (distance of < 20 genes) (Fig. S4). Since being identified by two independent approaches, the genetic loci are likely to be involved in the production. The proteins herein identified as unique to the producer strains include a sulfurase and glutathione S-transferase that could be involved in the biosynthesis of roseobacticides (Table 2). Sulfur is a key component of both TDA and roseobacticides (Wang *et al*., 2016) and glutathione was proposed to be involved in the TDA resistance mechanism by *P. inhibens* (Wilson *et al*., 2016).

**Table 2.** Unique orthologous proteins to **A)** the producers M6-4.2, 8-1, DSM 17395 and DSM 26640 and **B)** the roseobacticide non-producer 27-4.

|  | **Locus tag in**<br>**DSM 17395** | **Description** | **GO classification** |
| --- | --- | --- | --- |
| 1 | PGA1_RS19085 | hypothetical protein | - |
| 2 | PGA1_RS14175 | transposase | transposase activity, sequence-specific DNA binding, transposition, DNA-mediated |
| 3 | PGA1_RS03520 | hemolysin-type calcium-binding region | calcium ion binding, intein-mediated protein splicing |
| 4 | PGA1_RS04200 | hypothetical protein | integral component of membrane |
| 5 | PGA1_RS02970 | sulfurase | catalytic activity, molybdenum ion binding, pyridoxal phosphate binding, metabolic process |
| 6 | PGA1_RS03280 | peptidase M50 | integral component of membrane |
| 7 | PGA1_RS04870 | class A beta-lactamase;<br>EC:3.5.2.6 | beta-lactamase activity, beta-lactam antibiotic catabolic process, response to antibiotic |
| 8 | PGA1_RS06640 | glutathione S-transferase;<br>EC:2.5.1.18 | glutathione transferase activity, metabolic process |
| 9 | PGA1_RS09195 | hypothetical protein | - |
| 10 | PGA1_RS09205 | hypothetical protein | - |
| 11 | PGA1_RS09210 | Lj965 prophage replication | - |
| 12 | PGA1_RS10850 | hypothetical protein | - |
| 13 | PGA1_RS14180 | integrase | nucleic acid binding, DNA integration |
| 14 | PGA1_RS19900 | hypothetical protein | - |

**B) Unique orthologues to roseobacticide non-producer 27-4**
|  | **Locus tag in 27-4** | **Description** | **GO classification** |
| --- | --- | --- | --- |
| 1 | PhaeoP14_03815,<br>PhaeoP14_03816,<br>PhaeoP14_03912,<br>PhaeoP14_03913,<br>PhaeoP14_03930,<br>PhaeoP14_03931 | uncharacterized protein | - |
| 2 | PhaeoP14_03813,<br>PhaeoP14_03915,<br>PhaeoP14_03928 | RNA polymerase sigma factor SigK | DNA binding, transcription factor activity, sequence-specific DNA binding, sigma factor activity, DNA-templated transcription, initiation, regulation of transcription, DNA-templated |
| 3 | PhaeoP14_03814,<br>PhaeoP14_03914,<br>PhaeoP14_03929 | hypothetical protein | membrane, integral component of membrane, plasma membrane |
| 4 | PhaeoP14_03817,<br>PhaeoP14_03911,<br>PhaeoP14_03932 | peptide-methionine (R)-Sulfoxide reductase;<br>EC:1.8.4, EC:1.8.4.12 | Peptide-methionine (R)-sulfoxide reductase activity, response to oxidative stress, protein repair, oxidation-reduction process |
| 5 | PhaeoP14_03663,<br>PhaeoP14_03731 | hypothetical protein | - |

Four of the orthologous proteins unique to the non-producer 27-4 represent a transposable element (Fig. S5). This element appears three times in the genome of 27-4 on the plasmids F, E and C. It covers the majority of plasmid F (13 kb) and thus, the transposable element could have entered the strain via this plasmid. The transposon located on plasmid C disrupts a gene cluster that contains four genes encoding a transcriptional regulator, an endonuclease, an oxidoreductase and an aldehyde dehydrogenase, with the transposable element inserted in the gene of the oxidoreductase (Fig. S5). Using this bioinformatic approach, we can only speculate that disruption of this gene cluster could have led to loss of the ability to produce roseobacticides and further experimentation by e.g. site-directed deletion is required.

The transposable element itself contains seven genes encoding a transposase, a resolvase, a RNA polymerase sigma factor, an anti-sigma E protein, two secreted surface proteins, and a peptide methionine sulfoxide reductase. The peptide methionine sulfoxide reductase is also present in the non-roseobactide producing TDA-producer *Ruegeria* sp. strain TM1040 (Fig. S6), but not in other *Ruegeria* or *Pseudovibrio* strains, making it unlikely as a shared reason for the non-production of roseobacticides.

The ability to biosynthesize roseobacticides appears to have been developed in a common ancestor of *P. inhibens, P. gallaeciensis* and *P. piscinae* in contrast to TDA production. The TDA-producer *Phaeobacter, Ruegeria* and *Pseudovibrio* are not phylogenetic neighbours, thus this feature possibly ‘jumped’ by horizontal gene transfer between different *Roseobacter-group* species. Furthermore, the strains selected for the analysis herein were obtained from different locations and time points demonstrating how conserved the phenotype of roseobacticide production is within those species.

### Bioactivity of roseobacticides against microalgae

Ethyl acetate extracts of DSM 17395 and 27-4 and their corresponding TDA-negative mutants were prepared from cultures grown under roseobacticide-inducing conditions. Only the extract of DSM 17395 contained roseobacticides as determined by HPLC-HRMS (Fig. S7). The strains were grown under iron limited conditions, which generally diminishes the production of TDA (D’Alvise *et al*., 2016) and correspondingly TDA was not present in detectable quantities.

The algae were treated with a roseobacticide extract 10-fold diluted with respect to the original bacterial culture (i.e. for roseobacticide-containing extract, the bacterial strains were cultivated in 25 mL ½YTSS with *p-*coumaric acid. The culture was extracted twice with 1:1 ethylacectate, dried and resuspended in 2.5 mL methanol. 50 μl of this roseobacticide-containing extract was added to 5 mL of algal culture for the bioassay). Cell numbers were assessed over 9 days. Growth of the cryptophyte *R. salina* (Fig. 2A) and the diatom *T. pseudonana* (Fig. 2B) was only affected by the roseobacticide-containing extract of the DSM 17395 wildtype. While the cell numbers initially dropped, the algae had recovered until the end of the experiment. The chlorophyte *T. suecica* (Fig. 2C) was not affected by any extract. Roseobacticides might also have an effect on *T. suecica* at higher concentrations, however, this could not be evaluated due to non-availability of pure standards. Initially, no growth of the haptophyte *E. huxleyi* was observed in cultures supplemented with any of the four extracts at the given concentration (Fig. S8). However, when the extract was diluted to a 100-fold, an inhibitory effect of the DSM 17395 wildtype extract was observed (Fig. 2D). Growth of the eukaryotic organisms was unaffected by the addition of diluent (methanol) (Fig. 2).

**Figure 2.**
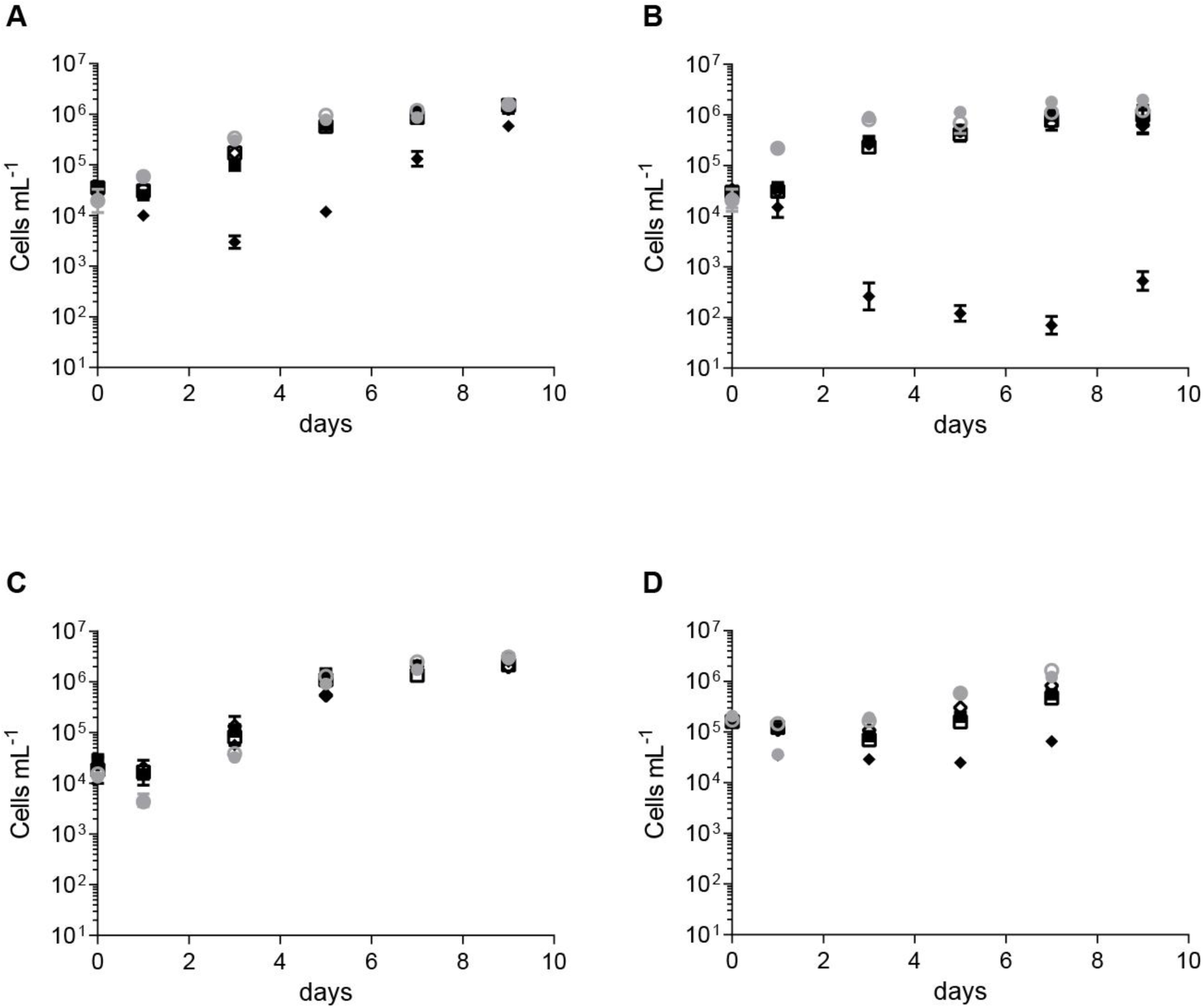
Bioactivity of *Phaeobacter* extracts against the microalgae A) *Rhodomonas salina*, B) *Thalassiosira pseudonana*, C) *Tetraselmis suecica* and D) *Emiliania huxleyi*. The extracts had a final concentration of 10% in A)-C) and 1% in D); percentage is given in regard to the original bacterial culture. ●: no addition, ◌: methanol, ♦:DSM 17395 wildtype, ◊: DSM 17395*tdaB*::Tn5, ■: 27–4 wildtype, □: 27–4*tdaB*::Tn*5*. Points are means of three replicates and error bars are standard deviations of the mean, except for D), which is one replicate.

In comparison to previous bioassays (Seyedsayamdost, Case, *et al*., 2011), we monitored microalgae representing four different groups (haptophyte, cryptophyte, heterokont, and chlorophyte) over nine days. We found the same impact on the cryptophyte *R. salina* and the haptophyte *E. huxleyi*. While the heterokont studied previously, *Chaetoceros muelleri*, was only weakly affected by roseobacticides, the heterokont *T. pseudonana* was highly affected in our experiments. The effect on any algae was temporary and the algae were able to recover, possibly due to the degradation of roseobacticides. Using the high concentration of *Phaeobacter* extracts, both the roseobacticide-containing and non-containing extracts had a high lytic effect on the haptophyte *E. huxleyi* (Fig. S8) suggesting that other so far undescribed metabolites play a role in the interaction with the *Roseobacter*-group bacteria.

### Bacterial-microalgal co-cultivation

The roseobacticide producer DSM 17395 or the non-producer 27-4 were co-cultivated with the microalgae *E. huxleyi* for 27 days and cell numbers were assessed (Fig. 3). The algae reached the maximum cell concentrations on days 12 *(E. huxleyi* + DSM 17395), 15 *(E. huxleyi* + 27-4) and 18 *(E. huxleyi)*. The algal cell concentrations of the axenic cultures compared to those in the co-cultivation setup with DSM 17395 were significantly lower on days 9 (*p*≤0.01) and 12 (*p*≤0.05) and significantly higher on days 21 (*p*≤0.0001), 24 (*p*≤0.001), 27 (*p*≤0.0001). There was no difference between the axenic cultures and those incubated with 27-4. Accordingly, co-cultivation setups of DSM 17395 and 27-4 differed in algal cell counts on day 12 (*p*≤0.05), 21 (*p*≤0.05), 24 (*p*≤0.05) and 27 (*p*≤0.01). The bacterial cell concentrations of both strains were significantly higher in the co-cultivation setups in comparison to the axenic samples of DSM 17395 and 27-4 from day 6 (*p*≤0.01) and 3 (*p*≤0.001) onwards, respectively. There was no significant difference of the bacterial numbers in the co-cultivations between DSM 17395 and 27-4, but on day 0 (*p*≤0.001).

**Figure 3.**
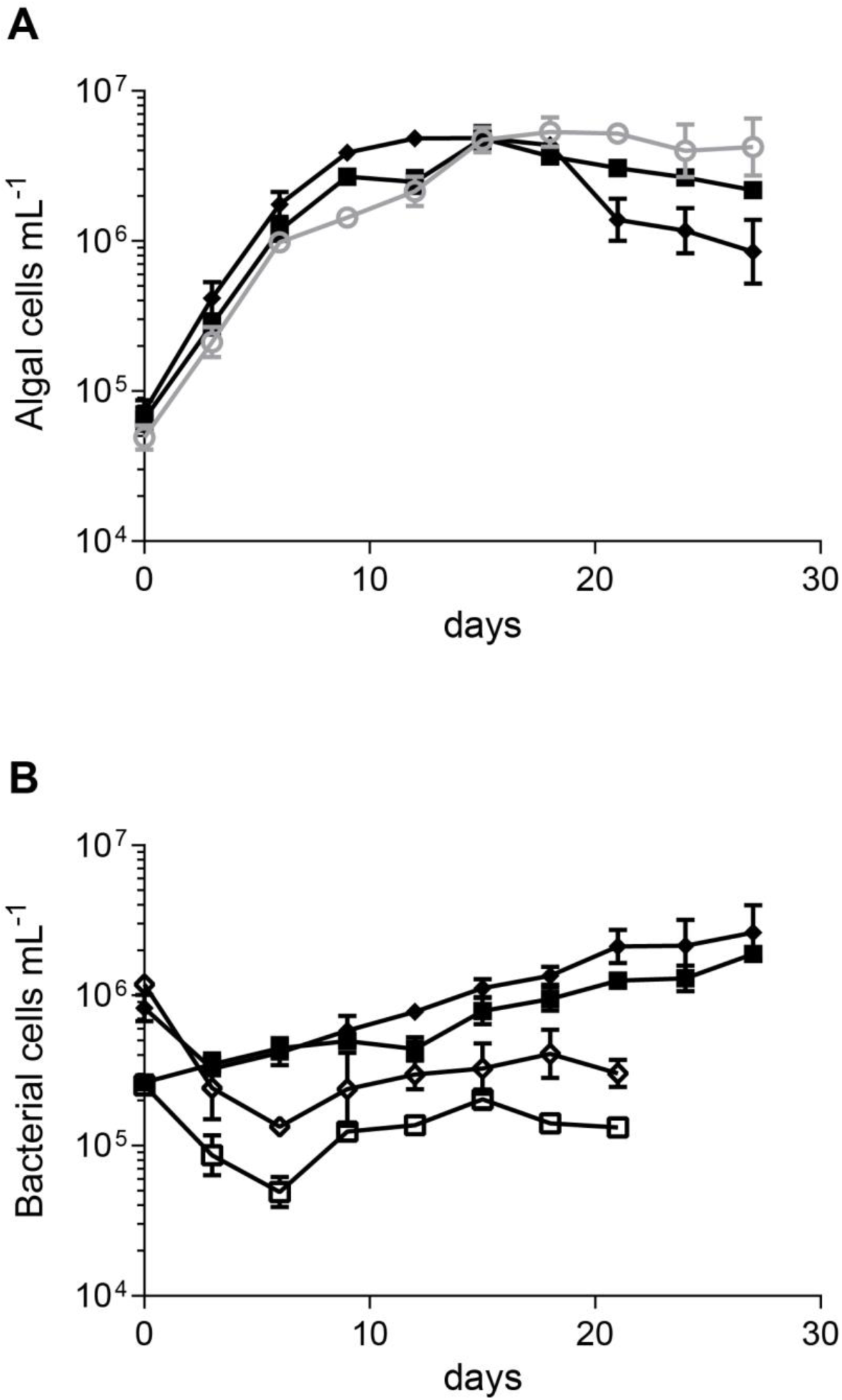
Co-cultivation of DSM 17395 wildtype and 27–4 wildtype with the microalgae *Emiliania huxleyi*. A) Algal cell counts, B) bacterial cell counts. Points are means of duplicates and error bars are standard deviations of the mean. ◌: *E. huxleyi*, ♦: *E. huxleyi* + DSM 17395, ■: *E. huxleyi* + 27–4, ◊: DSM 17395, □: 27-4.

Thus, we demonstrated the different direct effects on microalgae of two closely related TDA-producers; a roseobacticide producer and a non-producer. The roseobacticide producer promoted algal growth and enhanced their decay, an observation that was also found by a recent study on the metabolic dynamics of the interaction of *E. huxleyi* and *P. inhibens* (Segev *et al*., 2016b). While the enhanced decay might be attributed to the production of roseobacticides, the growth promotion would indicate that there is more to the beneficial effect of *P. inhibens* than TDA only. In contrast, the roseobacticide non-producer had no effect on the microalgae. The switch between mutualism and parasitism in algae-bacteria interactions has also been found in the interaction between the dinoflagellate *Prorocentrum minimum* and *Dinoroseobacter shibae* (Wang *et al*., 2014; H. Wang *et al*., 2015). It was suggested that *D. shibae* obtains its energy by degradation of polyhydroxyalkanoate and dimethylsulfoniopropionate (DMSP) (Wang *et al*., 2014), pathways that are also present in *P. inhibens* (Dickschat *et al*., 2010; Kanehisa *et al*., 2015) and with the latter molecule actually preventing the effect of TDA (Wichmann *et al*., 2016). Additionally, *E. huxleyi* is a known producer of DMSP (Wolfe and Steinke, 1996; Steinke *et al*., 2007). The growth promoting effect of bacterial indole-3-acetic acid demonstrated for the interaction between a diatom and *Sulfitobacter* appears unlikely due the inability of *Phaeobacter* to produce this specific molecule (Amin *et al*., 2015; Labeeuw *et al*., 2016). Furthermore, the establishment of these microbial interactions depend on motility, chemotaxis, attachment and quorum sensing (Sonnenschein *et al*., 2012; H. Wang *et al*., 2015), which are well-known characteristics of *Phaeobacter inhibens* (Newton *et al*., 2010; Berger *et al*., 2011; Gram *et al*., 2015; Wang *et al*., 2016). Thus, while some pathways appear universally important for microalgae-bacteria interactions, each species, and even each strain, might facilitate unique interactions. Such is the case in the herein presented phaeobacters where the interaction ranges from having no effect to detrimental effects on different microalgae.

### LC-HRMS and LC-MS/MS analysis of co-cultivation

Ethyl acetate extracts of upscaled DSM 17395 and *E. huxleyi* co-culture, collected at 17 and 27 days (early and late stationary phase of *E. huxleyi)*, were examined for the presence of TDA and roseobacticides by both full scan LC-HRMS (QTOF) and multiple reaction monitoring (MRM) LC-MS/MS (QqQ). Neither TDA or roseobacticides were detected in the co-culture at either time point.

In our experience, roseobacticides are produced in minute quantities, even in dense bacterial monocultures. Although roseobacticides were not detected in our LC-MS analysis in the co-cultures, it is plausible, if not likely, that roseobacticides are produced in quantities below our current limits of detection (LOD), even despite the approximately 5-10 fold lower LOD of the triple-quadrupole mass spectrometer (QqQ) compared to the QTOF.

## Conclusions

A mutualistic-parasitic interaction between the bacterium *P. inhibens* and the microalgae *E. huxleyi* has been proposed in previous studies (Seyedsayamdost, Case, *et al*., 2011; Segev *et al*., 2016a). This interaction is driven by the bacterial metabolites TDA and roseobacticides. It is not, however, known if all TDA producing roseobacters also produce roseobacticides. We here demonstrated that TDA production is a more widespread feature than roseobacticide biosynthesis and that the production of both compound groups is typical of the *Phaeobacter* genus. Furthermore, the proposed conversion from mutualistic to parasitic interaction of *P. inhibens* with the microalgae *E. huxleyi* was reproduced *in vitro*, while this effect was not observed in a closely related *Phaeobacter* strain that lost the ability of roseobacticide production.

The finding of a non-roseobacticide producing *Phaeobacter* strain, in combination with the fact that the microalgae used as aquaculture live feed were unaffected by the roseobacticides, is promising for future applications of phaeobacters as biocontrol agents in aquaculture. Also, roseobacticides are produced in very small quantities even when induced in bacterial culture and thus, *in situ* concentrations are presumably much lower than those tested in this study (not detectable with current techniques). Furthermore, we demonstrated that small genetic changes, possibly due to a transposable element, caused a different interaction phenotype with the ecologically important microalgae *E. huxleyi*, which thus may have larger environmental implications.

## Acknowledgements

*Rhodomonas salina* was kindly provided by Thomas Kiørboe (DTU Aqua, Denmark). *Phaeobacter porticola* P97^T^ was kindly provided by Sven Breider (University of Oldenburg, Germany). Thanks to Rasmus Kenneth Bojsen and Sven Anders Folkesson for help with flow cytometry (DTU Vet, Denmark). The collaboration with Boyke Bunk and Cathrin Spröer (DSMZ, Germany) on the genome sequences is acknowledged. Funding from the European Union (Seventh Framework Programs MaCuMBA (KBBE-2012-6-311975) and PharmaSea (KBBE-2012-6-312184)) and the Villum Kahn Rasmussen foundation (VKR023285) is acknowledged. Agilent Technologies is acknowledged for the Thought Leader Donation of the UHPLC-QTOF system.

## Supplementary information

Supplementary information containing materials and methods, supplementary tables and figures accompanies this article.

